# Glk1p is an actin fold metabolic enzyme whose polymerization is sensitive to nucleotide state

**DOI:** 10.64898/2026.07.09.731748

**Authors:** Michael D. Carver, Phillip Kyriakakis, Rachel M. Barry, Elena Monfort, Andres E. Leschziner, Mark A. Herzik, James E. Wilhelm

**Author notes:** Co-First Authors (Listed Alphabetically) (Michael D. Carver, Phillip Kyriakakis).

## Abstract

The actin fold is present in enzymes ranging from sugar kinases to chaperones. Previous work on the actin fold metabolic enzyme, glucokinase (Glk1p) in S. cerevisiae found it could form filaments in response to its substrates, ATP and glucose (Stoddard et al., 2020). Here, we have identified the product, glucose 6-phosphate (G6P), as a second trigger for Glk1p polymerization *in vitro*. Furthermore, the addition of ADP to G6P-Glk1p filaments causes filament disassembly, suggesting that polymerization is sensitive to the state of the bound nucleotide and/or the transfer of the gamma phosphate. We have also identified a specific metabolic state, the accumulation of G6P during stationary phase, that triggers Glk1p polymerization *in vivo*. While the structures of Glk1p filaments in either the ATP/glucose or G6P-bound form are not similar to conventional actin filaments, the sensitivity of assembly to the gamma phosphate of the nucleotide provides a conceptual bridge between the cytoskeleton and metabolic regulation via enzyme polymerization.

**Summary:** The yeast glucokinase, Glk1p, is an actin fold enzyme that undergoes regulated polymerization in response to ATP/glucose and glucose 6-phosphate (G6P). Elevated G6P during stationary phase triggers Glk1p filament assembly *in vivo* consistent with a role in regulating glycolytic flux.

## Introduction

The presence of a dynamic actin cytoskeleton was long considered one of the hallmarks of eukaryotic cells. However, the discovery of bacterial actins that form dynamic filaments erased this distinction between prokaryotes and eukaryotes (Jones et al., 2001; Møller-Jensen et al., 2002; Szwedziak et al., 2012). While both bacterial and cytoskeletal actins form a single clan within the actin superfamily (Stoddard et al., 2017), there is a wide variety of proteins that possess an actin fold. These actin fold enzymes have diverse functions such as protein folding (the Hsp70/DnaK family of chaperones), carbohydrate metabolism (hexo/glucokinases), and alarmone generation (PPX/GPPA phosphatases) (Bork et al., 1992; Flaherty et al., 1991; Kabsch and Holmes, 1995). Previous work on one of these proteins, the yeast glucokinase, Glk1p, found that the GFP-tagged protein formed filaments in log phase (Noree et al., 2019) and in response to glucose (Stoddard et al., 2020) *in vivo* and that the combination of glucose with ATP could trigger filament formation *in vitro* (Stoddard et al., 2020).

This suggested a connection between the actin cytoskeleton and the emerging area of metabolic enzymes that are regulated via polymerization. For instance, filament formation is intimately connected with the activation of Acetyl-CoA carboxylase, liver phosphofructokinase, and mammalian CTP synthetase (Beaty and Lane, 1983; Meredith and Lane, 1978; Webb et al., 2017; Lynch et al., 2017). In contrast, end product inhibition by CTP triggers the assembly of bacterial CTP synthetase filaments in an inactive conformation (Barry et al., 2014). Other work has shown enzymatic filament formation under stressful conditions such as low pH (Petrovska et al., 2014) or during developmental changes (Hugener et al., 2024). Furthermore, visual screens for metabolic enzyme structures suggest that this form of enzyme regulation is broadly distributed throughout metabolism (Noree et al., 2019). Thus, the further characterization of filament-forming actin fold enzymes outside of the clan of classical polymerizing actins could provide insights into both metabolic regulation and the routes available to evolving a dynamic cytoskeleton.

In this study, we have revisited the previous work on Glk1p in light of our observation that tagged Glk1p behaves differently from the native protein. We demonstrate that the yeast glucokinase, Glk1p, undergoes regulated polymerization *in vitro* in response to two distinct triggers: ATP+glucose or glucose 6-phosphate (G6P). Interestingly, ADP triggers the disassembly of G6P-Glk1p filaments, arguing that, like actin, Glk1p polymerization is sensitive to the state of the gamma phosphate of ATP. Furthermore, we have identified a metabolic state with elevated G6P that triggers the formation of Glk1p filaments *in vivo*. We propose that G6P-Glk1p filaments play a role in flux balancing between different parts of glycolysis when cells are in precarious metabolic states. Furthermore, the sensitivity of assembly to the state of the nucleotide suggests a possible evolutionary path for the development of dynamic filaments.

## Results and Discussion

### Glk1p forms filaments *in vitro* in response to substrate binding

In a previous screen of the 440 metabolic enzymes in the yeast GFP strain collection, we identified 60 metabolic enzymes that form structures during log phase, growth to saturation, or stationary phase (Noree et al., 2019). Interestingly, the only member of the actin superfamily identified in our screen was the yeast glucokinase, Glk1p-GFP, which forms foci in both log phase and stationary phase (Noree et al., 2019). Subsequent work by Stoddard et al. using a combination of *in vitro* biochemistry and GFP-tagged Glk1p argued that the trigger for Glk1p filament formation *in vivo* is ATP+glucose and that polymerization acts as a brake to prevent excess glucose from causing ATP levels to be depleted prior to ATP being generated by the later steps in glycolysis (Stoddard et al., 2020). While this model is attractive, we recently found that untagged Glk1p does not form visible filaments *in vivo* in response to glucose and that C-terminal tagging Glk1p drastically alters filament formation *in vivo* (Fig S1). Other polymerizing actin family proteins such as the bacterial actin MreB are also sensitive to tag identity and placement (Ouzounov et al., 2016; Swulius and Jensen, 2012), highlighting the need to investigate various tagging strategies and, when possible, native protein when studying filament dynamics. As a result, we explored the possibility that other types of regulators for Glk1p filament assembly might have been overlooked.

One of the most interesting features of Glk1p is that it is the first actin family member outside of the eucaryotic and bacterial cytoskeletal actins capable of filament formation (Stoddard et al., 2020). In order to explore and expand our understanding of the triggers for Glk1p filament formation, we adapted an actin pelleting assay for use with Glk1p (Wickline et al., 2016). While filament-forming actins only bind ATP, Glk1p has two substrates with the potential to regulate its assembly: glucose and ATP. Therefore, we tested whether glucose, ATP, or both glucose and ATP had any effect on the ability of Glk1p to form higher-order structures. Glk1p alone showed minimal pelleting when incubated with control buffer (3.4%), ATP alone (2.6%) or glucose alone (6.6%) (Fig. 1A). Thus, individual substrates are not sufficient to trigger assembly. In contrast, incubation of Glk1p with glucose and ATP caused 56.3% of the protein to pellet. Thus, the presence of both substrates is sufficient to cause Glk1p to form higher order structures (Fig. 1A). In contrast, fructose and galactose are not substrates of Glk1p and do not cause Glk1p pelleting in either the presence or absence of ATP (Fig. 1B). The Glk1p substrate, 2-deoxyglucose (Maitra, 1970), caused 43.5% of Glk1p to pellet in the presence of ATP and only 1.4% when it was the sole substrate in the reaction. Thus, the pelleting behavior of Glk1p exhibits substrate specificity. While these results together suggested that Glk1p was undergoing substrate-regulated polymerization, it was also possible that Glk1p was assembling into different types of structures depending on the trigger. To test this possibility, we examined the effects of ATP alone, glucose alone, ATP+glucose, and ATP+2-deoxyglucose on Glk1p by negative stain transmission electron microscopy (TEM) (Fig. 1C). Consistent with our pelleting assay, we did not observe any higher order structures when Glk1p was incubated with either ATP or glucose alone (Fig. 1C). In contrast, when the Glk1p assembly reaction contained both ATP and glucose, we observed the formation of extended Glk1p filaments ∼50-150nm in length (Fig. 1C). 2-deoxyglucose+ATP also triggered similar-sized Glk1p filament formation (Fig. 1C). This argues that Glk1p is capable of regulated filament formation in response to binding its substrates and/or the accumulation of its products, ADP and glucose 6-phosphate (G6P).

**Figure 1.**
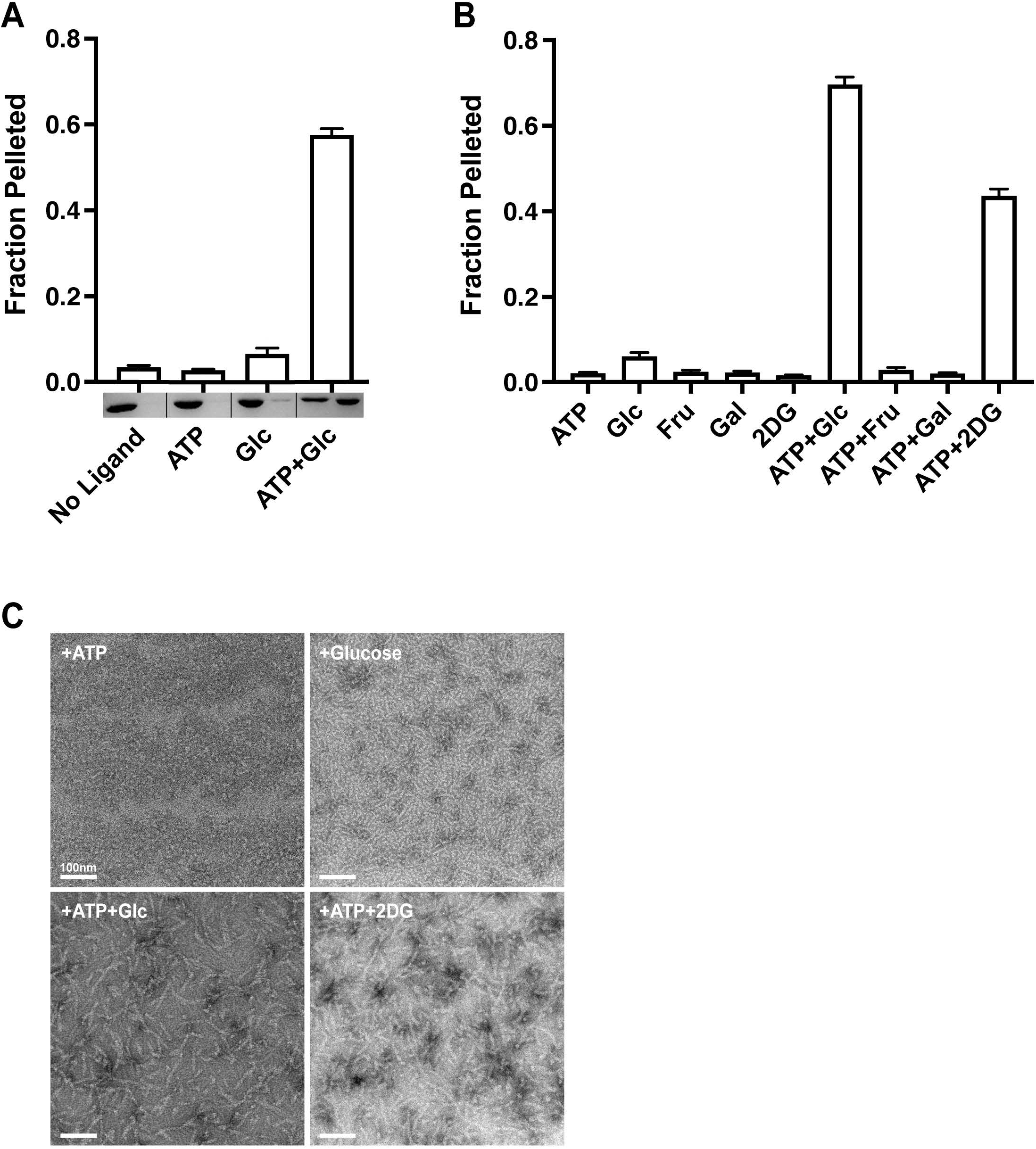
Glk1p forms filaments in response to its substrates. **(A**) Glk1p assembly was assayed by differential centrifugation. The average fraction of Glk1p pelleted is displayed ± SEM (n=6). Representative SDS-PAGE gel from the pelleting experiment is shown for each condition. The left band is the supernatant and the right band is the pellet for each condition. **(B)** Effect of different sugars on Glk1p pelleting. Average fraction of Glk1p pelleted is displayed ± SEM (n=6). **(C)** TEM images of Glk1p in buffer containing either 5mM ATP, 5mM glucose, 5mM ATP+glucose, or 5mM ATP+2-deoxyglucose. All scale bars are 100nm.

### Glk1p filament formation does not require ATP hydrolysis

While the assembly and disassembly of cytoskeletal actin is regulated by ATP hydrolysis, bacterial actins exhibit a diversity of behavior in response to ATP binding and hydrolysis (Bean and Amann, 2008; Derman et al., 2012; Garner et al., 2004; Singh et al., 2013). Therefore, in order to assess the relative roles of substrate binding and enzyme activity on Glk1p polymerization, we tested the effect of non-hydrolyzable analogs of ATP on Glk1p pelleting (Fig. 2A). While AMP-PNP alone caused little Glk1p pelleting (3.2%), assembly reactions with AMP-PNP+glucose caused 19.2% of the Glk1p to pellet (Fig. 2A). We observed similar effects with the non-hydrolyzable analog ATPγS: 2.5% of Glk1p pelleted with ATPγS alone, but 27.7% of Glk1p pelleted when both glucose and ATPγS were present in the reaction (Fig. 2A). These results suggest that ATP hydrolysis is not required for Glk1p filament formation – a conclusion that is supported by negative stain TEM of both the AMP-PNP+glucose and ATPγS+glucose assembly reactions (Fig. 2B). This argues that the binding of both substrates, ATP and glucose, is likely sufficient to trigger Glk1p assembly and ATP hydrolysis is not required.

**Figure 2.**
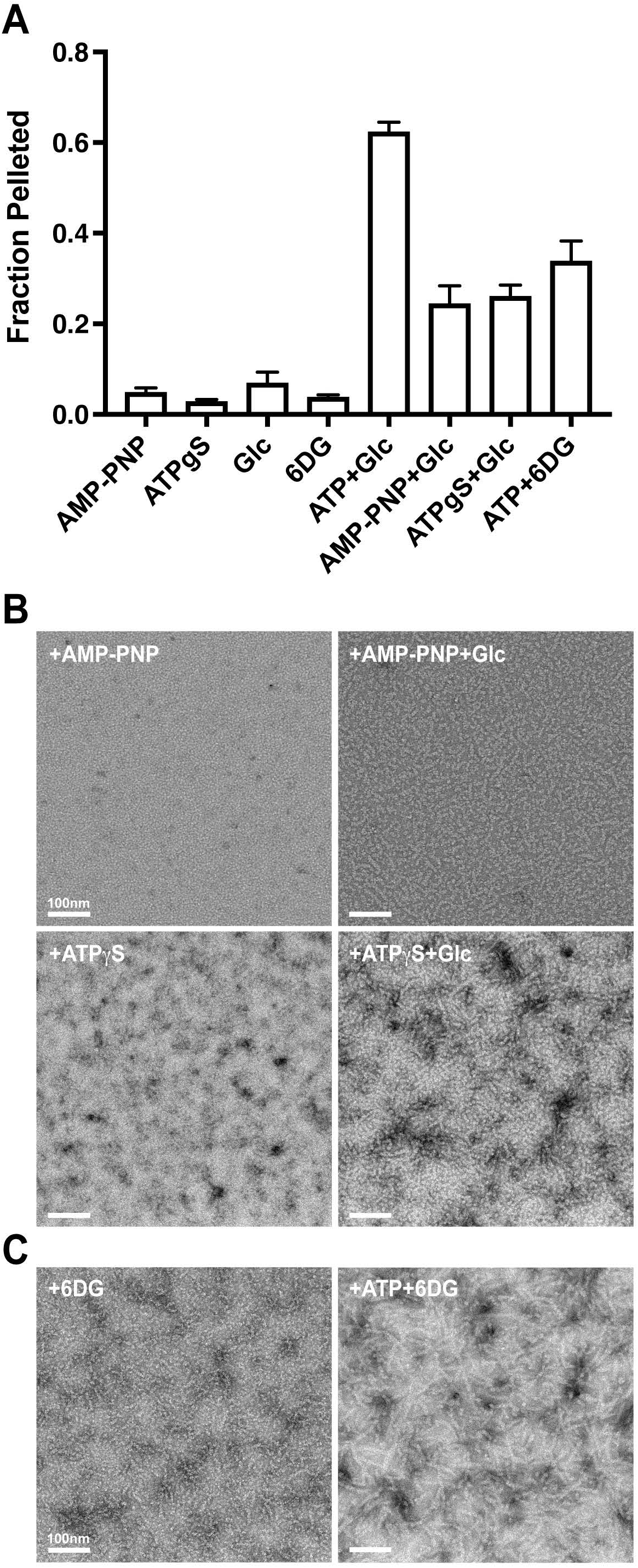
Glk1p filament formation does not require ATP hydrolysis. **(A)** The effect of non-hydrolyzable ATP analogs on Glk1p pelleting with and without glucose. The average fraction of Glk1p pelleted is displayed ± SEM (n=6). **(B)** TEM images of Glk1p diluted 1:5 in buffer containing either 5mM AMP-PNP alone, 5mM AMP-PNP with 5mM glucose, 5mM ATPγS alone, or 5mM ATPγS with 5mM glucose. **(C)** TEM images of Glk1p diluted 1:5 in buffer containing 5mM 6-Deoxyglucose alone or 5nM ATP with 5mM 6-Deoxyglucose. All scale bars are 100nm.

Since some enzymes are capable of hydrolyzing “non-hydrolyzable” analogs or distort the binding pocket of the enzyme, we also explored whether 6-Deoxyglucose, which can’t be phosphorylated, could trigger Glk1p filament formation in the presence of ATP. While 6-deoxyglucose alone caused little Glk1p pelleting (3.7%), assembly reactions with ATP +6-deoxyglucose caused 33.7% of the Glk1p to pellet in our assay (Fig. 2A). This pelleting result is supported by negative stain TEM of both the 6-deoxyglucose and the ATP+ 6-deoxyglucose assembly reactions (Fig. 2C).

### Glk1p polymerization is differentially regulated by each of its products, ADP and glucose 6-phosphate

While our results with non-hydrolyzable analogs of ATP and non-phosphorylatable 6-deoxyglucose argue that nucleotide hydrolysis is not required for Glk1p polymerization, they did not exclude the possibility that the products of the glucokinase reaction, ADP and glucose 6-phosphate, could independently regulate Glk1p filament formation. Interestingly, glucose 6-phosphate alone was sufficient to cause significant Glk1p pelleting (27.2%) in our assay while ADP alone caused only minimal pelleting (Fig. 3A). Furthermore, the Glk1p pelleting that we observe in the presence of glucose 6-phosphate is due to filament formation (Fig. 3A). The presence of both ADP+glucose-6-phosphate products in the assembly reaction, however, prevented Glk1p from pelleting (Fig. 3A). Thus, glucose 6-phosphate alone is sufficient to trigger Glk1p polymerization, while the addition of ADP is sufficient to inhibit the accumulation of G6P-induced Glk1p filaments.

**Figure 3.**
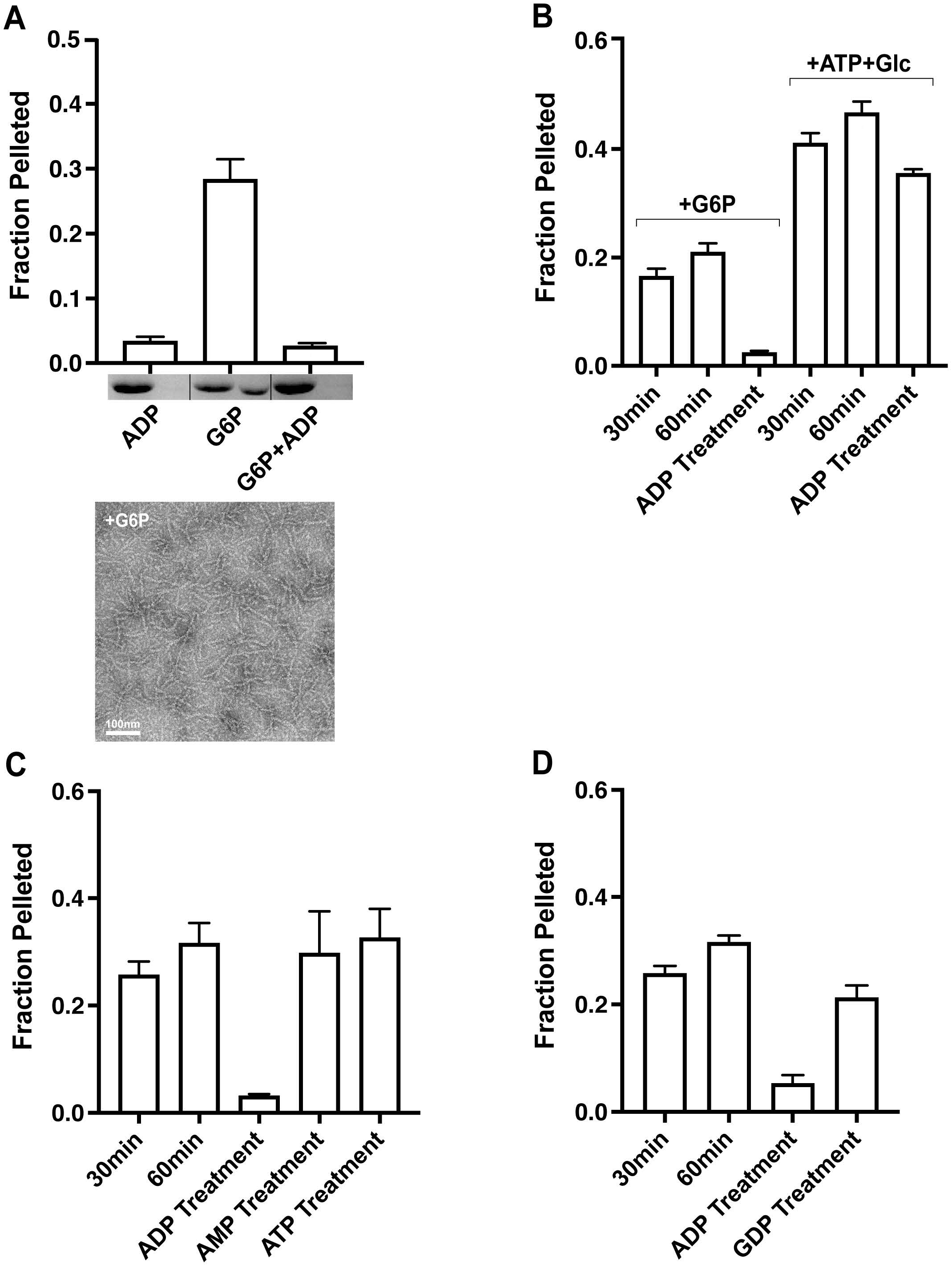
ADP promotes disassembly of G6P-Glk1p filaments. **(A)** The effect of ADP, glucose 6-phosphate (G6P), and ADP+G6P on pelleting. The average fraction of Glk1p pelleted is displayed ± SEM (n=6). Representative SDS-PAGE gel from pelleting experiment is shown for each condition. The left band is the supernatant and the right band is the pellet for each condition. Below is a TEM image of Glk1p in buffer containing 5mM G6P. **(B)** ADP added at the 30min time point of assembly reaction causes G6P-Glk1p filaments to disassemble but has limited effects on ATP/glucose-filaments. The effect of ADP was assayed 30 min after addition while control groups in G6P buffer were assayed for assembly at 30 min and 60 min time points. Assembly of Glk1p for each condition was measured using pelleting assay. The average fraction of Glk1p pelleted is displayed ± SEM (n=5). **(C-D)** The effect of ADP of disassembly of G6P-Glk1p filaments is specific for ADP when compared to **(C)** AMP and ATP or **(D)** GDP. The effect of each nucleotide was assayed 30 min after addition while control groups in G6P buffer were assayed for assembly at 30 min and 60 min time points as well. Assembly of Glk1p for each condition was measured using a pelleting assay. The average fraction of Glk1p pelleted is displayed ± SEM (n=4).

This result raised the question of whether or not ADP can promote the disassembly of existing filaments and whether the effects of ADP are specific to filaments formed in the presence of glucose 6-phosphate. In order to test the effects of ADP on pre-assembled Glk1p filaments, we incubated Glk1p for 30 min in the presence of the appropriate trigger (G6P or ATP+glucose) and then added ADP to the assembly reaction. The effects of ADP treatment on Glk1p filament formation were then assayed 30 min after addition using our pelleting assay. ADP treatment had only minimal effects on Glk1p filaments formed in the presence of ATP+glucose (Fig. 3B). In contrast, while Glk1p pelleting modestly increased between the 30 and 60 min time points in the glucose 6-phosphate reaction, addition of ADP at the 30 min time point triggered disassembly of Glk1p filaments (Fig. 3B). This effect is specific for ADP, since the addition of AMP, ATP, or GTP did not trigger disassembly (Fig. 3C-D). Thus, ADP can both block the assembly of G6P-Glk1p filaments and promote the disassembly of pre-existing G6P-Glk1p filaments, while only having weak effects on the disassembly of ATP+glucose filaments.

### Glk1p filaments formed in the presence of glucose 6-phosphate and ATP/glucose are structurally similar, but exhibit distinct polymerization behavior

The fact that glucose 6-phosphate and ATP/glucose could both trigger Glk1p filament formation, but had a differential response to ADP suggested that Glk1p could assemble into filaments that were structurally distinct. In order to address this question, we developed a 3D model of each type of filament. Negative stain electron-micrographs of Glk1p filaments assembled with either G6P or ATP/glucose were subjected to two successive rounds of 2D classification (Figure 4 A-D). The best particles in each dataset were then used to generate an *ab initio* 3D model of the two types of filaments (Figure 4 F-G). These models showed that Glk1p filaments assembled with either G6P or ATP/glucose showed a similar double-stranded antiparallel topology. Docking the model of the previously solved cryoEM structure (Stoddard et al., 2017) of ATP/glucose filaments into our G6P filament structure also supports that the interface and overall structure of the two types of Glk1p filaments are the same (Figure 4E)

**Figure 4.**
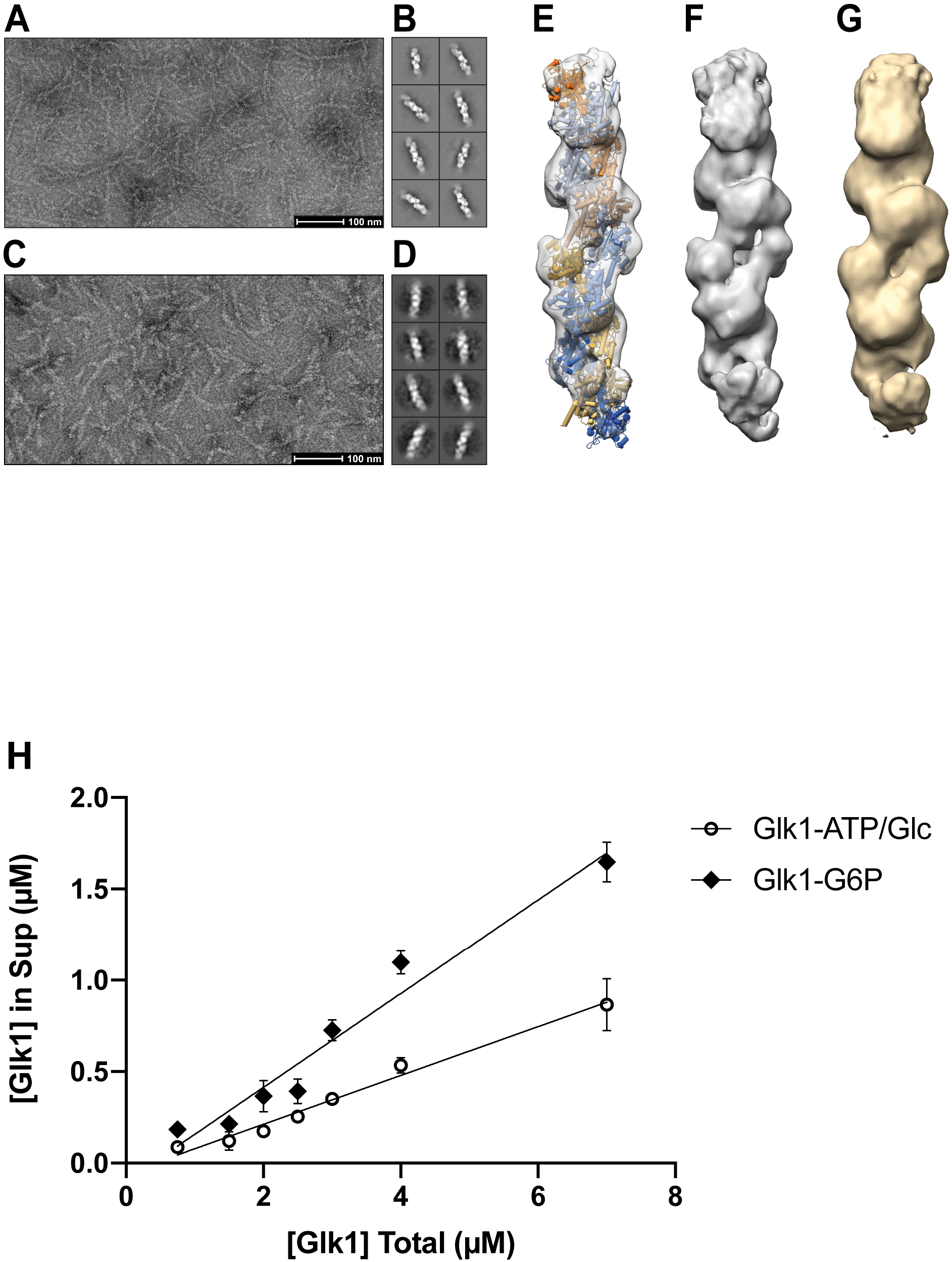
Glk1 is an antiparallel double helix which does not have a critical concentration. **(A-D)** Representative negative stain electron micrograph of Glk1p in the presence of (A) ATP/glucose or (C) glucose-6-phosphate (G6P) with representative 2D class averages (B and D, respectively). **(E-G)** (E) Glk1p+G6P shown in transparent gray surface with a docked model of the yeast Glk1p filament (adapted from PDB ID: 6PDT) colored by subunit, indicating that a similar overall anti-parallel double-stranded topology is shared by both Glk1p+ATP/glucose and Glk1p+G6P filament structures. Negative stain 3D reconstructions of Glk1p+G6P (F, gray) and Glk1p+ATP/glucose (G, wheat). **(H)** Proportion of Glk1p in the supernatant following differential centrifugation as a function of total Glk1p concentration, revealing no detectable critical concentration for Glk1p assembly in the presence of G6P (black triangles) or ATP/glucose (hollow circles) in the tested concentration range. The average fraction of Glk1p in the supernatant is displayed ± SEM (n=3).

We next examined whether the assembly behavior of Glk1p filaments differed when triggered by either G6P or ATP/glucose. In previous studies, Glk1p filaments assembled in the presence of ATP/glucose were found to have a critical concentration of ∼2uM (Stoddard et al., 2020). One of the hallmarks of cooperative assembly with a critical concentration is that increasing protein concentration above the critical concentration does not increase the amount of protein free in solution since the additional protein is incorporated into the polymer. In contrast, filaments that assemble isodesmically have no threshold for assembly, and the amount of unincorporated protein increases with increasing protein concentration (Skillman et al., 2013). Our measurements of the effect of increasing Glk1p concentration in the presence of ATP/glucose showed a linear increase in Glk1p free in solution. This argues that Glk1p assembled in the presence of ATP/glucose does not have a critical concentration and instead assembles isodesmically (Figure 4H). We also found that Glk1p in the presence of G6P also forms filaments that assemble isodesmically with no critical concentration (Figure 4H). Interestingly, while these two types of filaments are structurally similar, their assembly properties differ, with G6P being a less potent trigger of polymerization than ATP/glucose *in vitro* as measured by the fact that less Glk1p was in incorporated into filaments in the presence of G6P than ATP/glucose at every Glk1p concentration examined (Figure 4H). This argues that while the two classes of Glk1p filaments are structurally similar, they exhibit biochemical differences in their assembly that have implications for their function *in vivo*.

### Accumulation of glucose 6-phosphate triggers Glk1p filament formation *in vivo*

The fact that ATP/glucose and glucose 6-phosphate both promote Glk1p filament assembly raised the question of which trigger is most physiologically relevant. Our previous screen of the yeast GFP strain collection found that Glk1p-GFP formed puncta *in vivo* when shifted stationary phase growth to media with fresh glucose. Furthermore, Stoddard et al. found that Glk1p-GFP filaments form when yeast grown to saturation is shifted to fresh glucose (Stoddard et al., 2020). Together, these results suggested that Glk1p formed filaments in response to a large burst of glucose or glucose 6-phosphate. However, when we examined the effect of different C-terminal tags on Glk1p filament formation, we found conflicting results. C-terminal tagging of Glk1p with 3xHA allowed Glk1p to form filaments in response to shifting cultures to fresh glucose, while tagging with myc did not (Fig. S1). The conflicting results depending on the different C-terminal tags, GFP, 3xHA, and myc, argue that alterations to the C-terminus of Glk1p can have profound effects on Glk1p filament formation. As a result, we focused our efforts on examining the assembly behavior of untagged Glk1p *in vivo* using antibodies generated against Glk1p.

Immunofluorescence of native Glk1p did not reveal filament formation under growth conditions and/or media shifts that caused C-terminally tagged Glk1p to form filaments (Fig. S1). Since this failure to detect Glk1p filaments could have been due to a problem with our Glk1p antibody, we tested our Glk1p antibody by staining yeast strains where Glk1p was tagged with 3xHA which triggered filament formation. Our Glk1p antibody stained Glk1p-3xHA filaments and recognized no structures in a *GLK1Δ* strain, arguing that native Glk1p does not form large filaments under conditions that trigger Glk1p-GFP or Glk1p-3xHA to form filaments (Fig S2).

This raised the possibility that the metabolic state that triggers Glk1p filament formation is not commonly encountered during normal growth and/or the filaments that native Glk1p forms are below the level of detection in the absence of a tag that sensitizes the protein to filament formation. In order to explore this further, we examined the effects of Glk1p overexpression on polymerization. Interestingly, while overexpressed Glk1p does not form filaments in stationary phase yeast, it rapidly forms extensive filaments and sheets in 17.1% of cells when the yeast are shifted to media containing fresh glucose (Fig. S2). This suggests that native Glk1p can form regulated filaments *in vivo* when Glk1p levels are high enough.

We next explored the possibility that changes in the amount of the triggers we identified *in vitro* could cause native Glk1p to form filaments *in vivo* by generating strains where the genes that act downstream of Glk1p in glycolysis were deleted. If glucose 6-phosphate is a major trigger of Glk1p polymerization *in vivo*, we predicted that strains defective in the next step of glycolysis, phosphoglucoisomerase (*PGI1*), would accumulate glucose 6-phosphate and trigger Glk1p filament formation when exposed to glucose (Fig. 5A). Consistent with this prediction, we found that when yeast were grown to stationary phase and then shifted to media with fresh glucose (YPD), 25.6% of *PGI1Δ* cells had Glk1p filaments as compared to 7.2% in unshifted (Fig. 5B). Additionally, this effect is not due to changes in Glk1p expression level in the *PGI1Δ* strain (Fig. 5B) and can be rescued by integrating PGI1 into the HIS3 locus (Fig. 5C). While this result suggested that high levels of glucose 6-phosphate could be a trigger for Glk1p assembly, it was also possible that Glk1p filaments were forming due to other defects associated with a general block in glycolysis. The closer a block in glycolysis is to the step producing glucose 6-phosphate, the more glucose 6-phosphate should accumulate when the mutant strain is exposed to glucose. As a result, if glucose 6-phosphate is the main trigger for Glk1p polymerization *in vivo*, we would expect that deleting *PGI1* would have the strongest effect on Glk1p polymerization, while deleting genes that act later in glycolysis, such as fructose-1,6-bisphosphate aldolase (*FBA1*) and phosphofructokinase (*PGK1*), would cause fewer Glk1p filaments to form (Fig. 5A). Consistent with this prediction, we observed a distinct gradient in the effect of glycolytic mutants on Glk1p polymerization (Fig. 5B). Upon shifting stationary phase cultures to fresh YPD, *PGI1* deletion strains exhibited the most Glk1p filament formation (25.6% of cells), followed by *FBA1* (9.0% of cells), while deleting *PGK1*, the furthest step in glycolysis we examined, displayed no detectable Glk1p polymerization in response (Fig. 5B).

**Figure 5.**
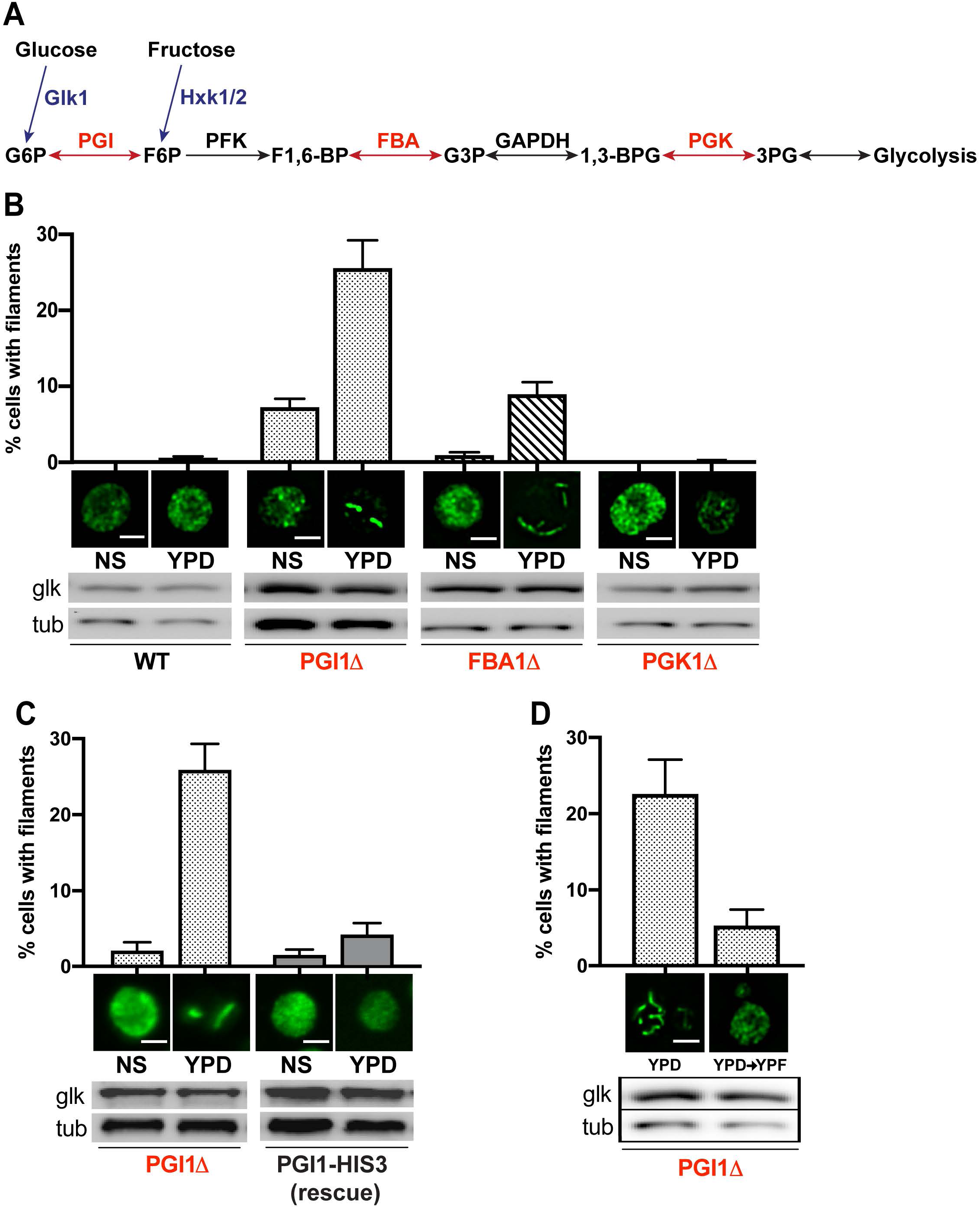
Blocks in glycolysis that cause G6P accumulation promote Glk1p filament formation. **(A)** The upper portion of the glycolytic pathway is shown for reference. Metabolites: G6P=glucose 6-phosphate, F6P=fructose 6-phosphate, F1,6-BP=fructose-1,6-bisphosphate, G3P=glyceraldehyde-3-phosphate, 1,3,BPG=1,3-bisphosphoglycerate, 3PG=3-phosphoglycerate. Enzymes: PGI=phosphoglucoisomerase, PFK=phosphofructokinase, FBA=fructose-1,6-bisphopshate aldolase, GAPDH=glyceraldehyde 3-phosphate dehydrogenase, PGK=phosphoglycerate kinase. Fructose enters glycolysis one step after where glucose enters. Genes shown in red were deleted and assayed for their effect on Glk1p filament formation. **(B)** Wild type and glycolysis mutants were grown for 5 days (stationary phase) and then shifted to YPD for 30 minutes. NS is strain prior to shift. YPD is strain 30 min after shift to YPD. The percentage of cells with Glk1p filaments was determined by immunofluorescence. The average percentage of cells with Glk1p filaments is displayed ± SEM (WT n=5; *PGIΔ* n=14; *FBA1Δ* n=6; *PGK1Δ* n=6). Immunoblots showing the level of Glk1 and tubulin proteins for each strain and condition are shown. All scale bars are 3μm. **(C)** The effect of deleting *PGI1* on Glk1p filament formation is rescued by integration of the *PGI1* ORF into the endogenous *HIS3* locus. The average percentage of cells with Glk1p filaments is displayed ± SEM (n=4) All scale bars are 3μm. **(D)** Shifting *PGI1Δ* strains to YPD for 30min triggers Glk1p filament formation, but a subsequent shift to YPF for 30min causes the filaments to disassemble. The average percentage of cells with Glk1p filaments is displayed ± SEM (n=4). All scale bars are 3μm.

These results argue that an accumulation of glucose 6-phosphate *in vivo* is sufficient to trigger Glk1p filament formation and that the effects that we observe are not merely due to disrupting glycolysis.

Together, these results suggest that Glk1p filaments form *in vivo* in response to a specific metabolic state: glucose 6-phosphate accumulation during stationary phase. However, it was unclear whether these filaments formed reversibly. Since fructose enters glycolysis immediately after the *PGI1* step, we reasoned that fresh fructose would promote exit from stationary phase and result in Glk1p disassembly (Fig. 5D). To test this, we triggered Glk1p polymerization in stationary phase *PGI1Δ* cells by shifting to fresh YPD. After 30 min, the cells were then shifted to YPF. Consistent with our prediction, 22.6% of the *PGI1Δ* cells exhibited Glk1p filaments when shifted to YPD, while only 5.3% of the cells had Glk1p filaments following a subsequent shift to YPF (Fig. 5D). This set of media shifts did not change in Glk1p protein levels, arguing that the absence of filaments is due to disassembly rather than protein degradation (Fig. 5D).

Thus, Glk1p filaments form in response to glucose 6-phosphate accumulation during stationary phase and these filaments subsequently disassemble upon successfully exiting stationary phase through a YPF-induced rescue of glycolysis.

## Discussion

The identification of an ever-increasing number of metabolic enzymes that can form filaments and other structures argues that the regulation of enzyme activity via polymerization is a widespread phenomenon (Chong et al., 2015; Mazumder et al., 2013; Narayanaswamy et al., 2009; Noree et al., 2010, 2019; O’Connell et al., 2014; Shen et al., 2016; Tkach et al., 2012). The ubiquity of metabolic enzyme structures also suggests that this mode of enzyme regulation could have been the basic substrate for the evolution of the classical cytoskeleton (Barry and Gitai, 2011). Consistent with this, previous work on Glk1p found that the GFP-tagged protein formed filaments *in vivo* during log phase (Noree et al., 2019) and in response to glucose (Stoddard et al., 2020). These previous studies also found that the combination of glucose with ATP could trigger filament formation *in vitro* (Stoddard et al., 2020). However, the cryoEM structure of the glucose+ATP Glk1p filaments revealed a filament structure and polymerization interface distinct from that used by conventional actins (Stoddard et al., 2020), arguing that the parallels between actin and Glk1p polymerization are limited.

Our finding that the assembly properties of Glk1p are significantly perturbed by a variety of tags led us to reexamine the triggers of *in vitro* and *in vivo* Glk1p assembly. While we found that Glk1p is triggered by glucose+ATP, we also found that glucose-6-phosphate alone is sufficient to trigger Glk1p polymerization. Surprisingly, ADP can trigger disassembly of G6P-Glk1p filaments (Figure 3B), arguing that the ability of Glk1p to polymerize is sensitive to the position of the gamma phosphate – a sensitivity that is reminiscent of the role of ATP hydrolysis in regulating actin filament dynamics (Korn et al., 1987). It is worth noting that while both actin and Glk1p are sensitive to the state of the nucleotide, the actual sequence of events is quite different. The release of the gamma phosphate triggers actin depolymerization, whereas the presence of both ADP and G6P promotes Glk1p disassembly.

While detailed structural studies of ATP/glucose-Glk1p filaments, G6P-Glk1p filaments, and the ADP/G6P-Glk1p monomer will be necessary to better understand how Glk1p substrates and products regulate polymerization, the extensive literature on hexokinases suggests a simple model for Glk1p polymerization. Hexokinases are induced-fit enzymes where glucose binds to the open state of the enzyme, triggering a conformational change to a closed state that sequesters the sugar within its binding site (Bennett and Steitz, 1978; Fuente et al., 1970; Steitz et al., 1981). The subsequent binding of ATP triggers a further conformational change to an enzymatically primed state, promoting the phosphotransfer reaction (Fuente et al., 1970; Steitz et al., 1981; Wilson and Schwab, 1996). Glucose 6-phosphate binding can also cause hexokinase to assume a closed conformation (McDonald R. C. et al., 1979). Our work suggests that the closed states of ATP/glucose-Glk1p and G6P-Glk1p are both competent for polymerization.

While little is known about the structure of the ADP/G6P-bound hexokinases, we propose that ADP/G6P-Glk1p assumes a partially open conformation that is incompatible with polymerization. This would explain why the addition of ADP causes G6P-assembled Glk1p filaments. One implication of this model is that if ATP/glucose-Glk1p filaments were enzymatically active, ATP/glucose-Glk1p filaments should display dynamic behavior similar to that of actin. Future work aimed at determining whether ATP/glucose-Glk1p filaments are enzymatically active and/or identifying mutations that make Glk1p active within the polymer should provide valuable insight into how a metabolic enzyme can be converted into an actin-like polymer.

Polymerization can be used to tune the allosteric regulation of an enzyme (Johnson and Kollman, 2020). While both ATP/glucose and glucose 6-phosphate are triggers for polymerization *in vitro*, we have only found evidence that glucose 6-phosphate drives filament formation *in vivo*. Furthermore, glucose 6-phosphate accumulation only triggers Glk1p polymerization during a specific metabolic state – stationary phase. The fact that filament assembly is restricted to stationary phase suggests that additional metabolic features of this cellular state, such as low levels of ADP, may contribute to regulating Glk1p polymerization *in vivo*. Consistent with this, while yeast maintain an energy charge ([ATP]+1/2[ADP]/[ATP+ADP+AMP]) of 0.8-0.9 as they transition from log phase to stationary phase (Ball and Atkinson, 1975), recent single-cell measurements of ATP levels have identified a low ATP subpopulation of yeast cells that emerges as glucose is depleted and becomes dominant as the culture progresses to stationary phase (Luzia et al., 2024). The constant energy charge combined with decreasing ATP levels implies that ADP levels must also decline in these cells. The drop in ADP levels in this stationary phase subpopulation could explain why *PGI* deletion strains, which block the entry of glucose 6-phosphate into glycolysis, only trigger Glk1p filament formation in stationary phase yeast. These findings open the door to future work on how filament assembly might interact with metabolic heterogeneity in yeast populations undergoing nutrient stress.

One attractive feature of regulating metabolic enzymes via polymerization is that it can allow novel forms of control of metabolic pathways. For instance, enzymes that assemble cooperatively into filaments have a critical concentration that results in the amount of free enzyme being constant when the total amount of enzyme is above the critical concentration. This can potentially buffer metabolic systems and control flux. In contrast, isodesmic polymerization has no critical concentration and has not been described for any class of metabolic filament.

Interestingly, while previous studies found that Glk1p assembles cooperatively with a critical concentration of ∼2μM, we have found no evidence of a critical concentration for Glk1p assembled in either the presence of ATP/glucose or G6P arguing that Glk1p can assemble isodesmically. While isodesmic assembly might seem at odds with Glk1p being an actin-related polymer, actin in *T. gondii* assembles isodesmically, arguing that cooperative assembly is not an essential feature of filament assembly in the actin superfamily (Skillman et al., 2013).

Furthermore, recent work on septins has shown that they can assemble both isodesmically and cooperatively, depending on reaction conditions and cofactors (Vogt et al., 2025). This raises the possibility that Glk1p has two modes of assembly that can be modulated *in vivo* to generate different types of metabolic control. Such dynamic control could be particularly valuable in supporting metabolic heterogeneity as yeast encounter changing nutrient environments. Future studies will be needed to shed light on how these modes of assembly are regulated *in vitro* and *in vivo*.

## Materials and Methods

### Cloning, expression, and purification of yeast Glk1p

#### Cloning

The full-length coding region of *GLK1* was subcloned into the pPROEX-HTc expression vector, generating an N-terminal 6xHis-tagged version of *GLK1* (*His_6x_-GLK1*) that was validated by sequencing.

#### Protein expression and purification

*His_6x_-GLK1* construct was transformed into Rosetta^TM^ Competent Cells (Novagen) and fresh transformants were inoculated into LB-Amp and grown overnight. The overnight culture was then diluted into 1L LB-amp to an initial OD_600_ of 0.05 and grown at 37^°^C to an OD_600_ of 0.2. The culture was then shifted to 30^°^C and grown to an OD_600_ of 0.5. IPTG (Sigma) was then added to the culture at a final concentration of 200 mM to induce protein expression for 14-16 hours at 30^°^C. Cells were then pelleted, resuspended in lysis buffer (50mM Tris pH 8.0, 20mM imidazole, 500mM NaCl, 0.5mM TCEP and 1x Protease Inhibitor Cocktail (Sigma)), and stored at −80^°^C until purification.

Frozen *His_6x_-GLK1* Rosetta^TM^ cells were quickly thawed. Fresh protease inhibitors were added during the thawing procedure. All subsequent purification procedures were conducted at 4^°^C. Cells were lysed via six rounds of 30-second sonication (550 Sonic Dismembrator, ThermoFischer). Following lysis, the lysate was centrifuged at 29,000g for 30 min and the supernatant was incubated with 2ml HisPure^TM^ Ni-NTA Resin (ThermoFisher) for 4 hours. The lysate-bead mixture was then batch-washed twice with 50ml wash buffer (lysis buffer without protease inhibitor), packed into a column, and washed with an additional 20 bed volumes. His_6x_-Glk1p was eluted into fifteen 1ml fractions with elution buffer (wash buffer supplemented with 250mM imidazole). Peak protein containing fractions were pooled and the protein concentration was measured by a Bradford Protein Assay Kit (Bio-Rad). TurboTEV protease (Eton Biosciences) was then added at a 50:1 protein to protease ratio to the pooled His_6x_-Glk1p and incubated overnight during dialysis against 20mM Tris pH 8.0, 500mM NaCl, and 0.5mM TCEP(dialysis buffer). Following digestion, the sample was recovered from the dialysis membrane and incubated with Ni-NTA resin in dialysis buffer for 4 h at 4 °C. The resin was then batch washed twice with dialysis buffer and transferred to a column. The initial flow-through was collected and the resin was washed with 10 mL dialysis buffer lacking imidazole. Under these conditions, cleaved Glk1p remained weakly associated with the Ni-NTA resin and was not recovered with the 0 mM imidazole wash. Cleaved Glk1p was eluted with 20 mM imidazole in dialysis buffer and collected in ten 1 mL fractions, whereas His₆-Glk1p was eluted with 200 mM imidazole in dialysis buffer to assess cleavage efficiency. Peak protein-containing fractions, as determined by Ponceau staining, were pooled and concentrated to 1 ml using an Amicon® Ultra-4 Centrifugal Filter Unit. Gel filtration was then performed using an ÄKTA™ pure protein purification system in 20mM Tris pH 8.0, 500mM NaCl, 0.5mM TCEP and 10% glycerol in order to remove the imidazole and prepare the sample for storage at −80^°^C. Fractions containing untagged Glk1p were then pooled, aliquoted, flash frozen, and stored at −80^°^C until the protein was needed. Enzymatic activity of purified Glk1p was assayed by continuous spectrophotometric assay following previous protocols in order to confirm that the protein was functional (EC 2.7.1.2 Sigma).

#### Glk1p Pelleting Assay

Glk1p samples were thawed and prespun at 285,000g for 30 min/30^°^C and the protein concentration of the supernatant was assayed via Bradford as previously described. All reactions were fully assembled and equilibrated to 30^°^C prior to addition of Glk1p with a final reaction volume of 100μl. All pelleting reactions were carried out using 5.5mM Glk1p, 20mM Tris pH 8.0, and 55mM NaCl. All products or substrates in the reaction (ATP, ADP, AMP, GDP, AMP-PNP, ATPγS, MgCl_2_, glucose, fructose, galactose, 2-deoxyglucose, glucose 6-phosphate) were used at a final concentration of 5mM. Reactions were then incubated at 30^°^C for 30 min and then spun at 218,000g for 30 min/30^°^C to pellet Glk1p structures. Following this final spin, the supernatant and pellet were separated, denatured, and stored with SDS-PAGE sample buffer (SB) (100μl 1x SB to resuspend pellet, 50μl 2x SB + 50μl supernatant).

The supernatant and pellet of each reaction was analyzed by SDS-PAGE (15ul supernatant and 7.5μl pellet samples). Gels were stained with Coomassie Brilliant Blue Dye (ThermoFisher) for 25 min and destained O/N before imaging on a FluorChem^TM^ using a 0.5 second exposure. FIJI image analysis was then used to quantify gel band area and intensity using the program’s specified protocol (imagej.nih.gov, Gels Submenu).

#### Disassembly assays

For assays involving disassembly of Glk1p filaments, 300μl reactions were assembled as above and incubated at 30^°^C for 30 min. The reaction was then separated into three equal sub reactions: 30 min control reactions were immediately spun at 218,000g/30^°^C for 30 min and samples were collected; 60 min control reactions were left untreated, incubated for another 30 min at 30^°^C, and spun at 218,000g/30^°^C for 30 min and samples were collected; “AMP/ADP/ATP/GDP treatment” reactions were treated with specified nucleotide (5mM of the additional nucleotide and 5mM MgCl_2_) at the 30 min mark immediately following splitting the master reaction and then treated and analyzed identically to the 60 min control fractions.

### Negative-Stain TEM Imaging and Data Processing

100μl Glk1p assembly reactions were assembled and incubated identically to the Glk1p pelleting assay before grid application. Samples for NSTEM imaging were prepared by applying 4µl of 305 μg/ml (5.5μM) Glk1p from each assembly reaction to glow discharged amorphous carbon grids which were then stained with 2% (w/v) uranyl acetate following established protocols (Scarff et al., 2018). Grids were examined and imaged using a FEI Tecnai Sphera microscope (FEI co.) operating at 200 kV fitted with a Gatan US4000 4k x 4k CCD camera for the following assembly conditions: (ATP, ATP+glucose, AMP-PNP, AMP-PNP+glucose). For all other conditions, grids were prepared and imaged with the help of the Core Microscopy Facility at Scripps Research using an 80 KV Talos L120C microscope (ThermoFisher) fitted with a CETA 16M CMOS camera.

Filamentous Glk1p particles were identified in cryoSPARC v3 (Punjani et al., 2017) using an elliptical blob (20 Å x 100 Å) before extraction at a 128-pixel box size (3.56 Å/pixel) for subsequent 2-D classification and sorting. Classes containing clear filamentous Glk1p were then subjected to *ab initio* model generation for subsequent homogenous refinement using gold-standard FSC estimation. Final maps were used without modification. PDB: 6PDT was rigid body fit into the corresponding EM density and expanded based on filamentous architecture.

### Yeast strains and media

All yeast strains used in this study were derived from *MAT-a his3Δ1 leu2Δ0 met15Δ0 ura3Δ0 (S288C)*. All yeast strains were grown at 30^°^C in YPD with the following exceptions: *PGK1Δ* and *FBA1Δ* mutants were grown in liquid culture media containing 0.2%glucose+YP+2%ethanol+2%glycerol+3%acetate or liquid and solid media containing YP+2%ethanol+2%glycerol+3%acetate. *PGI1Δ* mutants were grown in YP+2% fructose.

### Plasmid and DNA methods

For plasmid yeast transformation, yeast genomic tagging of specific loci, or yeast gene disruption, the LiOAc method was used (Ito et al., 1983). Genomic tagging and gene disruption were accomplished by transforming yeast strains with a PCR product that encoded the G418 resistance cassette, kanMX4. The DNA for this transformation was generated in one of two ways. For *PGI1Δ* strains, 5’ and 3’ 50-bp flanking sequences homologous to the gene of interest were synthesized into primers amplifying the antibiotic resistance cassette (Baudin et al., 1993; Brachmann et al., 1998; Longtine et al., 1998). For *FBA1Δ*:kanMX4 and *PGI1Δ*:kanMX4 disruption strains, diploid genomic DNA from the yeast Heterozygous Diploid A Complete Set (Invitrogen cat. Num. 95401.H4R3) was digested to inhibit PCR amplification of the wild-type copy. This allowed for amplification of knockout cassettes directly from the digestion reaction with longer 5’ and 3’ flanks homologous to the gene of interest than the oligosynthesis method. Flanks of 300bp or larger were designed with this second method. Cells transformed with the antibiotic cassette were allowed to grow on YPD or YPF for ∼24 hours and then replica plated onto plates containing carbon sources metabolizable by the desired knockout strain, along with the appropriate antibiotic. G418 was used at a final concentration of 400mg/ml. Gene disruption or genomic tagging was confirmed by PCR. All PCR was performed with KOD hot start polymerase (EMD) in 50μl reactions (1x KOD buffer, 1.25mM MgSO_4_, 200mM dNTP, 0.3mM of each oligonucleotide, and 10-20ng of template) using the manufacturer’s protocols. The *GLK1*-linker-3xHA strain was generated by tagging *GLK1* at the endogenous locus using pFA6a-3HA-Kan plasmid and 50bp oligos that targeted the 3’ end of the *GLK1* gene.

Glk1p with different C-terminal tags were made by first amplifying the *GLK1*-linker-3xHA from genomic DNA from the yeast made with pFA6a-3HA-Kan into a pFA6a-3HA with SAL1-HF and PAC1 sites added to PCR primers. Next, a 3’ oligo containing a copy of the desired tag (linker-3xHA and linker-MYC), a stop codon, and PAC1 site was used to amplify *GLK1* and replace the sequence on the pFA6a-*GLK1*-linker-3xHA construct with the new tag.

Dual expression plasmid pCEV-G4-Km was purchased from Addgene #46819 (Vickers et al., 2013). A hygromycin version of the plasmid (pCEV-G4-Hygro) was generated by performing a marker swap technique (Kelly and Hoffman, 2002; Muhlrad et al., 1992). *GLK1* overexpression was achieved by cloning *GLK1* into pCEV-G4-Hygro using Spe1 and Pac1 restriction sites. This put *GLK1* under the control of the constitutive *HXT7* promoter.

*PGI1Δ* strains were rescued by integrating the PGI1 ORF into the HIS3 locus. The *PGI1* coding DNA sequence plus 500bp of its 5’UTR and 3’UTR was cloned XhoI/NotI into pRS403. 5’ and 3’ 50-bp flanking sequences homologous to HIS3 were synthesized into primers amplifying the *PGI1-HIS3* cassette (*5’UTR-PGI1 ORF-PGI1 3’UTR-HIS3*). Deletion strains were transformed with this cassette, and transformants were selected by growing on SC His^-^ + G418. Successful integration of *PGI1* into the *HIS3* locus was confirmed via PCR.

### Media shift experiments

*FBA1Δ* and *PGK1Δ* strains were grown in YP+2%ethanol+2%glycerol+3%acetate overnight, then back-diluted into 0.2%glucose+YP+2%ethanol+2%glycerol+3%acetate and grown to stationary phase (5 days). *PGK1Δ* strains were grown in YP + 2% fructose overnight, then back-diluted into fresh YPF and grown to stationary phase (5 days). Stationary phase cultures were then shifted to test media (YP, YP + 2% glucose (YPD), or YP + 2% fructose (YPF)) for 30min at 22^°^C unless otherwise noted. In order to shift to test media, 4.5ODs of cells were pelleted, rinsed once with test media, and resuspended into 600ml of the test media (YP, YP + 2% glucose (YPD), or YP + 2% fructose (YPF)). Non-shifted cells were pelleted (4.5 OD_600_) and resuspended into 600ml of non-shifted media. Cells were scored as the percentage of cells with filaments compared with total cell counts.

### Antibody generation

The full-length coding region of Glk1p was cloned into pPROEX-HTc C containing an N-terminal 6xHis-tag and expressed in BL21 *E coli.* Cells were lysed in buffer containing 50mM phosphate pH 7.4, 250mM NaCl, 2mM Mg^2+^, 10mM imidazole, fresh 1.5mg/100ml DNaseI (Sigma Cat#D4527), fresh 1.5mg/100ml RNase A (Qiagen Mat#1007885), fresh 1mM BME, and fresh protease inhibitor cocktail (Sigma Cat#P8340). After resuspension in lysis buffer, cells were processed using a Laboratory Microfluidizer® (Microfluidics Corporation model M-110S). Soluble Glk1p was column purified using Ni-NTA affinity beads (Qiagen), eluted in wash buffer containing 250mM imidazole, dialyzed in 100mM PO_4_ buffer, pH 7.0, and injected into rabbits (antiserum production by Covance).

### Immunostaining

Two OD_600_ of cells were fixed with 3.7% formaldehyde for 1 hour, washed in SK buffer (1M sorbitol, 45mM K_2_HPO_4_, and 7mM KH_2_PO_4_), and treated with zymolase (Zymo Research) for cell wall digestion. Next, cells were attached to Poly-D-Lysine-coated multiwell slides and permeabilized in MeOH and acetone for 5 minutes and 30 seconds, respectively. Blocking was in PBS + 2% BSA for 10 minutes. Primary antibodies used in this study were anti-Myc (1:200, 9E10, Invitrogen), anti-HA (1:500, 12CA5, SCBT), and anti-Glk1p (1:2000) and were diluted in PBS + 2% BSA and used to incubate at 4^°^C overnight. Next, cells were rinsed 2 times in PBS + 2% BSA and washed 3 times for 10 minutes in PBS + 2% BSA. Alexa488-anti-mouse (Invitrogen #A11029) and Alexa-568-anti-rabbit (Invitrogen #A11011) secondary antibodies were diluted in PBS + 2% BSA (1:200) and incubated for at least 1.5 hours. Cells were rinsed 2 times in PBS +2% BSA, DAPI (2mg/ml) stained for 10 minutes in PBS +2% BSA, rinsed once, and washed once in PBS + 2% BSA. Finally, cells were washed in PBS for 10 minutes before the slides were mounted with Vectashield® (Vector Laboratories #H-1000) and then sealed.

### Microscopy

Microscopy was performed using a DeltaVision Restoration Microscopy System (Applied Precision) and microscope (IX70; Olympus) using software SoftWoRx (Applied Precision). Quantitated images were taken with the same microscope and exposure settings as the controls from each experiment using a 60X 1.4NA Olympus PlanApo objective. For overexpression experiments, the longest exposure that would not saturate the camera was taken for cell counts and were kept the same for all images in each experiment. These settings filtered out the lowest expressing cells and the remaining cells scored for Glk1p structures.

### Protein sample preparation and Western blot analysis

Either 2 or 4 OD_600_ of cells were pelleted, and protein was extracted using an alkaline lysis method adapted from (Kushnirov, 2000). Briefly, cells were lysed with 0.1N NaOH for 5 minutes at room temperature, spun at 10,000’g and then resuspended in SDS lysis buffer containing a protease inhibitor cocktail (Sigma Cat# P8340). Protein was resolved with 10% SDS-PAGE and then transferred onto nitrocellulose membranes (BioRad) using the Owl-HEP1 electroblotting system (Thermo Scientific). Blots were probed with anti-Glk1p antibodies generated as described above at 1:2000 or anti-tubulin at 1:5000 (12G10, DSHB).

## Acknowledgements

We thank Scott Henderson, Theresa Fassel, and Kimberly Vanderpool of the Core Microscopy Facility at The Scripps Research Institute for their assistance in helpful discussions, sample preparation, and image collection. Work from the Wilhelm lab was supported by a grant from the HCIA program of HHMI. The yeast parent strain and yeast heterozygous knockout collection strains used in this study were a gift from L. Pillus (UC San Diego, La Jolla, CA). The yeast homozygous knockout collection strain was a gift from M. Niwa (MAT-a). The vector used to construct G418 deletion strains (pRH728) was a gift from R. Hampton (UC San Diego).

## Author Contributions

This work was devised and planned by M. Carver, P. Kyriakakis, A. Leschziner, M. Herzik, and J. Wilhelm. Experiments were carried out by M. Carver, P. Kyriakakis, R. Barry, and E. Monfort. Data were analyzed by M. Carver, P. Kyriakakis, M. Herzik, and E. Monfort. The manuscript was written by M. Carver, P. Kyriakakis, and J. Wilhelm with input from all authors.

---

**Supplementary Figure 1.**
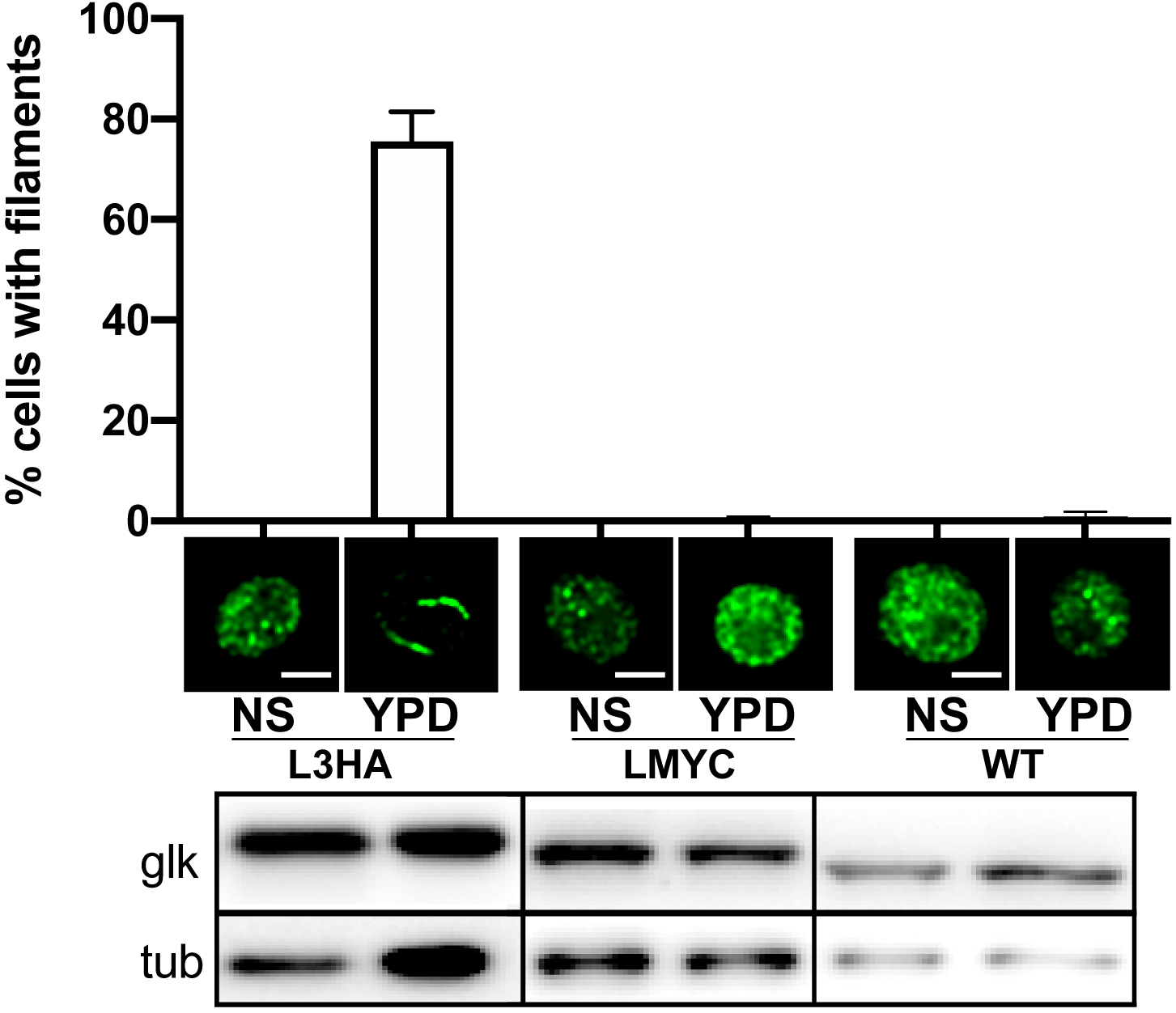
Tagging the C-terminus of Glk1p changes its polymerization behavior *in vivo* Wild type yeast and strains where *GLK1* was tagged at the endogenous locus with a C-terminal tag were grown for 16 hours then shifted into YPD for 30 minutes. The percentage of cells with Glk1p filaments was then assayed by immunofluorescence. The average percentage of cells with Glk1p filaments is displayed ± SEM (n=3). Representative images with either anti-HA, anti-myc, or anti-Glk1p are shown. All scale bars are 3μm. Immunoblots showing the level of each Glk1p and tubulin are shown.

**Supplementary Figure 2.**
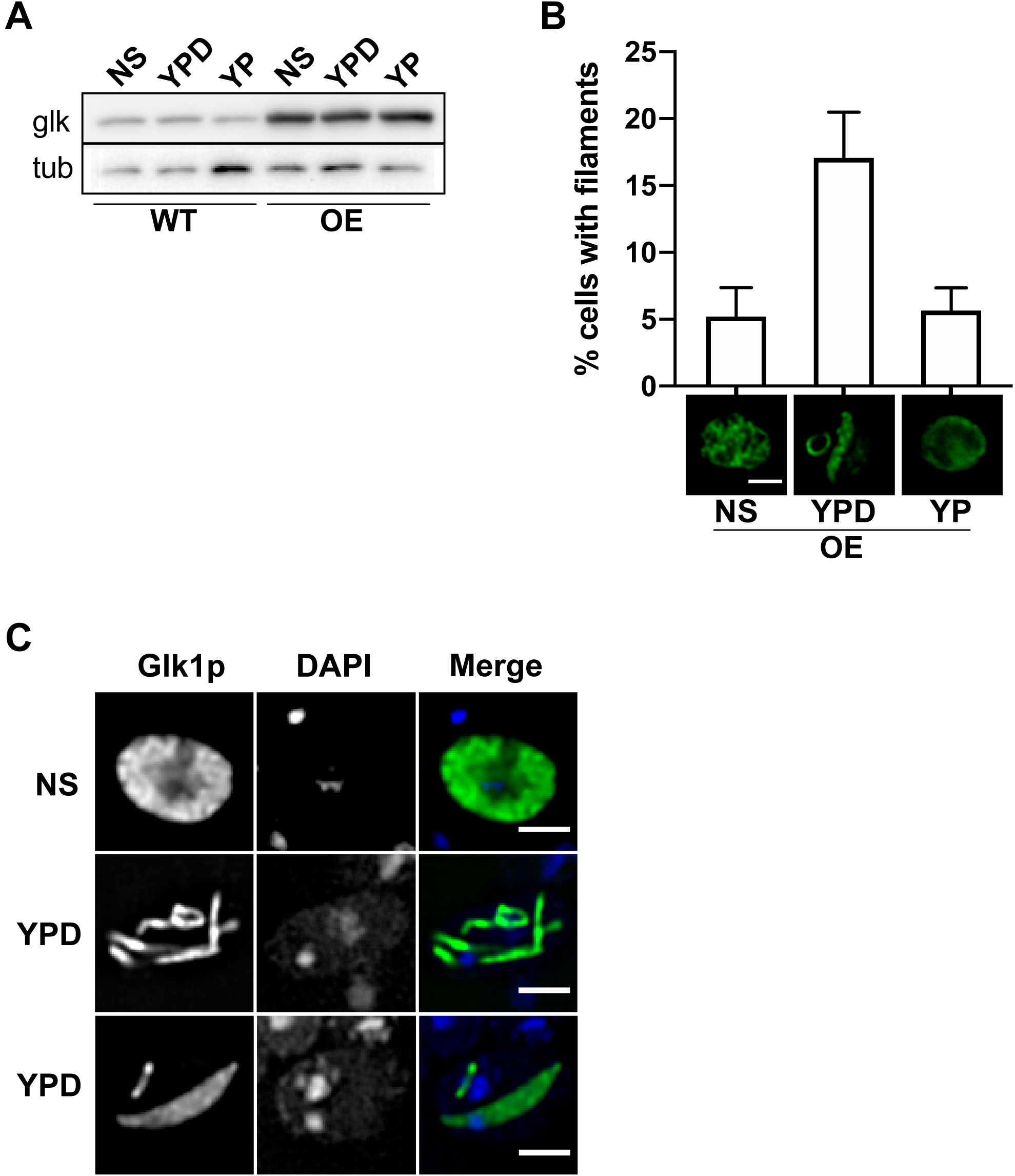
Overexpressed Glk1p polymerizes *in vivo* in response to the addition of fresh glucose. **(A)** Untagged Glk1p was overexpressed from a 2μm plasmid. Immunoblots of Glk1p and tubulin protein levels when strains overexpressing Glk1p (OE) or wild type (WT) are grown for 24 hours and then shifted to YPD or YP for 30 min. NS represents samples prior to shift and YPD or YP after the corresponding shift. **(B)** Percentage of cells with Glk1p filaments when strains overexpressing untagged Glk1p are assayed by immunofluorescence under the experimental conditions described in (A). The average percentage of cells with Glk1p filaments is displayed ± SEM (n=22) **(C)** Additional images of Glk1p structures that form when strains overexpressing Glk1p are grown for 24 hours and then shifted to YPD for 30 min. NS represents samples prior to shift where Glk1p is disbursed. After shift to YPD Glk1p forms filaments (middle row) or sheets (bottom row).

## References

Ball, W.J., and D.E. Atkinson. 1975. Adenylate energy charge in Saccharomyces cerevisiae during starvation. J Bacteriol. 121:975–982. doi:10.1128/jb.121.3.975-982.1975.

Barry, R., and Z. Gitai. 2011. Self-assembling enzymes and the origins of the cytoskeleton. Curr Opin Microbiol. 14:704–711. doi:10.1016/j.mib.2011.09.015.

Barry, R.M., A.-F. Bitbol, A. Lorestani, E.J. Charles, C.H. Habrian, J.M. Hansen, H.-J. Li, E.P. Baldwin, N.S. Wingreen, J.M. Kollman, and Z. Gitai. 2014. Large-scale filament formation inhibits the activity of CTP synthetase. eLife. 3. doi:10.7554/eLife.03638.

Baudin, A., O. Ozier-Kalogeropoulos, A. Denouel, F. Lacroute, and C. Cullin. 1993. A simple and efficient method for direct gene deletion in Saccharomyces cerevisiae. Nucleic Acids Res. 21:3329–3330.

Bean, G.J., and K.J. Amann. 2008. Polymerization Properties of the T. Maritima Actin, Mreb. Biochemistry. 47:826–835. doi:10.1021/bi701538e.

Beaty, N.B., and M.D. Lane. 1983. Kinetics of activation of acetyl-CoA carboxylase by citrate. Relationship to the rate of polymerization of the enzyme. J. Biol. Chem. 258:13043–13050.

Bennett, W.S., and T.A. Steitz. 1978. Glucose-induced conformational change in yeast hexokinase. Proc Natl Acad Sci U S A. 75:4848–4852.

Bork, P., C. Sander, and A. Valencia. 1992. An ATPase domain common to prokaryotic cell cycle proteins, sugar kinases, actin, and hsp70 heat shock proteins. Proc Natl Acad Sci U S A. 89:7290–7294.

Brachmann, C.B., A. Davies, G.J. Cost, E. Caputo, J. Li, P. Hieter, and J.D. Boeke. 1998. Designer deletion strains derived from Saccharomyces cerevisiae S288C: a useful set of strains and plasmids for PCR-mediated gene disruption and other applications. Yeast. 14:115–132. doi:10.1002/(SICI)1097-0061(19980130)14:2%3C115::AID-YEA204%3E3.0.CO;2-2.

Chong, Y.T., J.L.Y. Koh, H. Friesen, S.K. Duffy, K. Duffy, M.J. Cox, A. Moses, J. Moffat, C. Boone, and B.J. Andrews. 2015. Yeast Proteome Dynamics from Single Cell Imaging and Automated Analysis. Cell. 161:1413–1424. doi:10.1016/j.cell.2015.04.051.

Derman, A.I., P. Nonejuie, B.C. Michel, B.D. Truong, A. Fujioka, M.L. Erb, and J. Pogliano. 2012. Alp7R Regulates Expression of the Actin-Like Protein Alp7A in Bacillus subtilis. J Bacteriol. 194:2715–2724. doi:10.1128/JB.06550-11.

Flaherty, K.M., D.B. McKay, W. Kabsch, and K.C. Holmes. 1991. Similarity of the three-dimensional structures of actin and the ATPase fragment of a 70-kDa heat shock cognate protein. Proc Natl Acad Sci U S A. 88:5041–5045.

Fuente, G.D., R. Lagunas, and A. Sols. 1970. Induced Fit in Yeast Hexokinase. European Journal of Biochemistry. 16:226–233. doi:10.1111/j.1432-1033.1970.tb01075.x.

Garner, E.C., C.S. Campbell, and R.D. Mullins. 2004. Dynamic Instability in a DNA-Segregating Prokaryotic Actin Homolog. Science. 306:1021–1025. doi:10.1126/science.1101313.

Hugener, J., J. Xu, R. Wettstein, L. Ioannidi, D. Velikov, F. Wollweber, A. Henggeler, J. Matos, and M. Pilhofer. 2024. FilamentID reveals the composition and function of metabolic enzyme polymers during gametogenesis. Cell. 187:3303–3318.e18. doi:10.1016/j.cell.2024.04.026.

Ito, H., Y. Fukuda, K. Murata, and A. Kimura. 1983. Transformation of intact yeast cells treated with alkali cations. J Bacteriol. 153:163–168.

Johnson, M.C., and J.M. Kollman. 2020. Cryo-EM structures demonstrate human IMPDH2 filament assembly tunes allosteric regulation. Elife. 9. doi:10.7554/eLife.53243.

Jones, L.J.F., R. Carballido-López, and J. Errington. 2001. Control of Cell Shape in Bacteria: Helical, Actin-like Filaments in Bacillus subtilis. Cell. 104:913–922. doi:10.1016/S0092-8674(01)00287-2.

Kabsch, W., and K.C. Holmes. 1995. The actin fold. The FASEB Journal. 9:167–174. doi:10.1096/fasebj.9.2.7781919.

Kelly, D.A., and C.S. Hoffman. 2002. Gap repair transformation in fission yeast to exchange plasmid-selectable markers. Biotechniques. 33:978, 980, 982. doi:10.2144/02335bm02.

Korn, E.D., M.F. Carlier, and D. Pantaloni. 1987. Actin polymerization and ATP hydrolysis. Science. 238:638–644. doi:10.1126/science.3672117.

Kushnirov, V.V. 2000. Rapid and reliable protein extraction from yeast. Yeast. 16:857–860. doi:10.1002/1097-0061(20000630)16:9%3C857::AID-YEA561%3E3.0.CO;2-B.

Longtine, M.S., A.M. Iii, D.J. Demarini, N.G. Shah, A. Wach, A. Brachat, P. Philippsen, and J.R. Pringle. 1998. Additional modules for versatile and economical PCR-based gene deletion and modification in Saccharomyces cerevisiae. Yeast. 14:953–961. doi:10.1002/(SICI)1097-0061(199807)14:10%3C953::AID-YEA293%3E3.0.CO;2-U.

Luzia, L., J. Battjes, E. Zwering, D. Jansen, C. Melkonian, and B. Teusink. 2024. A fast method to distinguish between fermentative and respiratory metabolisms in single yeast cells. iScience. 27:108767. doi:10.1016/j.isci.2023.108767.

Lynch, E.M., D.R. Hicks, M. Shepherd, J.A. Endrizzi, A. Maker, J.M. Hansen, R.M. Barry, Z. Gitai, E.P. Baldwin, and J.M. Kollman. 2017. Human CTP synthase filament structure reveals the active enzyme conformation. Nat Struct Mol Biol. 24:507–514. doi:10.1038/nsmb.3407.

Maitra, P.K. 1970. A Glucokinase from Saccharomyces cerevisiae. J. Biol. Chem. 245:2423–2431.

Mazumder, A., L.Q. Pesudo, S. McRee, M. Bathe, and L.D. Samson. 2013. Genome-wide single-cell-level screen for protein abundance and localization changes in response to DNA damage in S. cerevisiae. Nucleic Acids Res. 41:9310–9324. doi:10.1093/nar/gkt715.

McDonald R. C., T.A. Steitz, and D.M. Engelman. 1979. Yeast hexokinase in solution exhibits a large conformational change upon binding glucose or glucose 6-phosphate. Biochemistry. 18:338–342. doi:10.1021/bi00569a017.

Meredith, M.J., and M.D. Lane. 1978. Acetyl-CoA carboxylase. Evidence for polymeric filament to protomer transition in the intact avian liver cell. J. Biol. Chem. 253:3381–3383.

Møller-Jensen, J., R.B. Jensen, J. Löwe, and K. Gerdes. 2002. Prokaryotic DNA segregation by an actin-like filament. EMBO J. 21:3119–3127. doi:10.1093/emboj/cdf320.

Muhlrad, D., R. Hunter, and R. Parker. 1992. A rapid method for localized mutagenesis of yeast genes. Yeast. 8:79–82. doi:10.1002/yea.320080202.

Narayanaswamy, R., M. Levy, M. Tsechansky, G.M. Stovall, J.D. O’Connell, J. Mirrielees, A.D. Ellington, and E.M. Marcotte. 2009. Widespread reorganization of metabolic enzymes into reversible assemblies upon nutrient starvation. Proc Natl Acad Sci U S A. 106:10147–10152. doi:10.1073/pnas.0812771106.

Noree, C., K. Begovich, D. Samilo, R. Broyer, E. Monfort, and J.E. Wilhelm. 2019. A quantitative screen for metabolic enzyme structures reveals patterns of assembly across the yeast metabolic network. Mol Biol Cell. 30:2721–2736. doi:10.1091/mbc.E19-04-0224.

Noree, C., B.K. Sato, R.M. Broyer, and J.E. Wilhelm. 2010. Identification of novel filament-forming proteins in Saccharomyces cerevisiae and Drosophila melanogaster. J. Cell Biol. 190:541–551. doi:10.1083/jcb.201003001.

O’Connell, J.D., M. Tsechansky, A. Royal, D.R. Boutz, A.D. Ellington, and E.M. Marcotte. 2014. A proteomic survey of widespread protein aggregation in yeast. Mol Biosyst. 10:851–861. doi:10.1039/c3mb70508k.

Ouzounov, N., J.P. Nguyen, B.P. Bratton, D. Jacobowitz, Z. Gitai, and J.W. Shaevitz. 2016. MreB Orientation Correlates with Cell Diameter in Escherichia coli. Biophys J. 111:1035–1043. doi:10.1016/j.bpj.2016.07.017.

Petrovska, I., E. Nüske, M.C. Munder, G. Kulasegaran, L. Malinovska, S. Kroschwald, D. Richter, K. Fahmy, K. Gibson, J.-M. Verbavatz, and S. Alberti. 2014. Filament formation by metabolic enzymes is a specific adaptation to an advanced state of cellular starvation. Elife. 3:e02409. doi:10.7554/eLife.02409.

Punjani, A., J.L. Rubinstein, D.J. Fleet, and M.A. Brubaker. 2017. cryoSPARC: algorithms for rapid unsupervised cryo-EM structure determination. Nat Methods. 14:290–296. doi:10.1038/nmeth.4169.

Scarff, C.A., M.J.G. Fuller, R.F. Thompson, and M.G. Iadanza. 2018. Variations on Negative Stain Electron Microscopy Methods: Tools for Tackling Challenging Systems. J Vis Exp. 57199. doi:10.3791/57199.

Shen, Q.-J., H. Kassim, Y. Huang, H. Li, J. Zhang, G. Li, P.-Y. Wang, J. Yan, F. Ye, and J.-L. Liu. 2016. Filamentation of Metabolic Enzymes in Saccharomyces cerevisiae. J Genet Genomics. 43:393–404. doi:10.1016/j.jgg.2016.03.008.

Singh, P., R.D. Makde, S. Ghosh, J. Asthana, V. Kumar, and D. Panda. 2013. Assembly of Bacillus subtilis FtsA: Effects of pH, ionic strength and nucleotides on FtsA assembly. International Journal of Biological Macromolecules. 52:170–176. doi:10.1016/j.ijbiomac.2012.09.019.

Skillman, K.M., C.I. Ma, D.H. Fremont, K. Diraviyam, J.A. Cooper, D. Sept, and L.D. Sibley. 2013. The unusual dynamics of parasite actin result from isodesmic polymerization. Nat Commun. 4:2285. doi:10.1038/ncomms3285.

Steitz, T.A., M. Shoham, W.S. Bennett, D.C. Phillips, C.C.F. Blake, and H.C. Watson. 1981. Structural dynamics of yeast hexokinase during catalysis. *Philosophical Transactions of the Royal Society of London. B*, Biological Sciences. 293:43–52. doi:10.1098/rstb.1981.0058.

Stoddard, P.R., E.M. Lynch, D.P. Farrell, A.M. Dosey, F. DiMaio, T.A. Williams, J.M. Kollman, A.W. Murray, and E.C. Garner. 2020. Polymerization in the actin ATPase clan regulates hexokinase activity in yeast. Science. 367:1039–1042. doi:10.1126/science.aay5359.

Stoddard, P.R., T.A. Williams, E. Garner, and B. Baum. 2017. Evolution of polymer formation within the actin superfamily. Molecular Biology of the Cell. 28:2461–2469. doi:10.1091/mbc.e15-11-0778.

Swulius, M.T., and G.J. Jensen. 2012. The helical MreB cytoskeleton in Escherichia coli MC1000/pLE7 is an artifact of the N-Terminal yellow fluorescent protein tag. J Bacteriol. 194:6382–6386. doi:10.1128/JB.00505-12.

Szwedziak, P., Q. Wang, S.M. Freund, and J. Löwe. 2012. FtsA forms actin-like protofilaments. EMBO J. 31:2249–2260. doi:10.1038/emboj.2012.76.

Tkach, J.M., A. Yimit, A.Y. Lee, M. Riffle, M. Costanzo, D. Jaschob, J.A. Hendry, J. Ou, J. Moffat, C. Boone, T.N. Davis, C. Nislow, and G.W. Brown. 2012. Dissecting DNA damage response pathways by analysing protein localization and abundance changes during DNA replication stress. Nat. Cell Biol. 14:966–976. doi:10.1038/ncb2549.

Vogt, E.J.D., I. Seim, W.T. Snead, B.N. Curtis, and A.S. Gladfelter. 2025. Cooperativity in septin polymerization is tunable by ionic strength and membrane adsorption. Biophys J. S0006-3495(25)00498–9. doi:10.1016/j.bpj.2025.08.005.

Webb, B.A., A.M. Dosey, T. Wittmann, J.M. Kollman, and D.L. Barber. 2017. The glycolytic enzyme phosphofructokinase-1 assembles into filaments. J Cell Biol. 216:2305–2313. doi:10.1083/jcb.201701084.

Wickline, E.D., I.W. Dale, C.D. Merkel, J.A. Heier, D.B. Stolz, and A.V. Kwiatkowski. 2016. αT-Catenin Is a Constitutive Actin-binding α-Catenin That Directly Couples the Cadherin·Catenin Complex to Actin Filaments. J Biol Chem. 291:15687–15699. doi:10.1074/jbc.M116.735423.

Wilson, J.E., and D.A. Schwab. 1996. Functional interaction of hexokinase with ATP requires participation by both small and large lobes of the enzyme: implications for other proteins using the actin fold as a nucleotide binding motif. The FASEB Journal. 10:799–801. doi:10.1096/fasebj.10.7.8635698.

